# An immunologist, ecologist and clinician walk into Plato’s cave to discuss infections

**DOI:** 10.1101/2025.03.04.641508

**Authors:** Yael Lebel, Avni S. Gupta, Victoria Chevée, Uri Alon, David S. Schneider

## Abstract

If the immune system is an interconnected network, then the evolution of an archetype that is ideal for fighting one pathogen should result in tradeoffs decreasing its ability to fight others. How many archetypes are there in an immune system? We infected diverse mice with *Plasmodium chabaudi,* and identified five distinct archetypes of responses based on the host’s position in microbial load, immune activity, and host damage space. To better understand the nature of these archetypes, we developed a mathematical model of a generalized host-pathogen system. This model explains the number, and distribution of archetypes across a population of diverse hosts. Mice resilient to *P. chabaudi* exhibited poor outcomes when challenged with influenza, SARS-CoV-1, or *Mycobacterium tuberculosis*, and vice versa, supporting our tradeoff hypothesis.

## Introduction

Plato’s allegory of the cave helps explain what we have been missing as we try to describe “disease space”^1^. Plato imagined prisoners in a cave who observed the world as a two-dimensional shadow on the cave’s wall. They are unable to reconstruct or comprehend the true three-dimensional (3D) structure of the world. Imagine there were three different rooms in Plato’s cave, each at ninety degrees to the next. Each room would present a different 2D view of the world to the inhabitants. If the prisoners could compare what they saw, together they could reconstruct a 3D reality. This is what we need to do with infectious disease research. Three independent models are commonly used to explain the within-host outcome of infectious diseases. First is the ecological concept of “disease tolerance” which describes health outcomes in relation to pathogen load ^2,3^. The second is the “damage response framework” which explains clinical health outcomes in relation to the intensity of the immune response ^4,5^. The third is the inflammatory response which records the mechanistic and functional immune reaction to an elicitor. Each field is self-contained and doesn’t inform the others; like the prisoners in the three caves, each of these approaches observes one of the three possible 2D views of a common 3D microbe-by-immunity-by-damage space (M x I x D) and cannot explain the whole system. Disease tolerance looks at microbes-by-damage (MxD), the damage response framework plots immunity-by-damage (IxD), and the third approach examines microbes-by-immunity (MxI). We can link ecological, clinical, and immunological thinking by examining the space occupied by different individuals responding to the same pathogen in MID space.

Complex systems face tradeoffs when they are tuned to a particular task; becoming magnificent at one function necessitates doing less well at another. Pareto described the outcome of this sort of multi-objective optimization a century ago for economic systems using a method that produces simple maps. Individuals who are the best at solving a task and lie at the vertices of a graph measuring the properties of a system are called archetypes^6–8^. The optimal solutions for the system lie between the archetypes. We can use the distribution of individuals on the multi-dimensional space describing the system to learn about the different unique tasks the system has evolved to excel in, and about the tradeoffs between these tasks^9^. For example, in an immune response, we could learn how the balance of the host’s relative differential investment in microbe killing versus repair of damage affects outcomes. How many archetypes should we expect to find in a simple system where microbes induce the immune response, the immune response clears microbes, and both the immune response and microbes may cause damage? What set of dimensions should we use to measure the shape of disease space? Here we show there are a limited number of immune archetypes in infected mice; we suggest this could result from multi-objective optimization and the dynamics of the immune response. We used diversity outbred mice (DO mice) and their 8 parent lines as a diverse population^10^. In plotting the results of more than 500 infections, we found that data point clouds of three types of damage (anemia, weight loss, and temperature loss) plotted against parasite load, and parasite clearance were roughly triangular. We mapped the phenotypes of the 8 parent lines in response to other infections onto these triangles and found that the mice that did well with malaria did poorly with TB, influenza, and SARS-COV1. To better understand how the dynamics of the immune response and repair could lead to these archetypes, we developed a mathematical model of a generalized host-pathogen system. This model characterizes the number, shapes, and distribution of response archetypes across a population of diverse hosts, integrating concepts from disease tolerance, the damage-response framework, and the inflammatory response into a unified description of infections within a microbe-immunity-damage space.

## Results

Like all animals, the mouse (Mus musculus) evolved its immune response under the selective pressure of multiple pathogens. We anticipated that signs of this evolution would persist in the many mouse strains collected by scientists and mouse fanciers. We used the shape these data points occupied in disease space to understand the multi-objective optimization of the immune response.

To measure the infection response of the mouse as a species, we used diverse mice generated by the collaborative cross. In this cross, eight strains of mice covering 95% of laboratory mouse diversity were intercrossed to produce stable lines^10^. These lines have themselves been intercrossed and then the offspring have been serially intercrossed to generate “Diversity Outbred (DO)” mice to create individual diverse mice^11^. To capture most of the timeline for acute infection and disease, we conducted a longitudinal study challenging DO mice with the murine malaria-causing agent *P. chabaudi* and monitoring them daily post-infection. We used a dose of *P. chabaudi* AJ that kills approximately 20% of infected C57BL/6 mice, and thus we anticipated finding strains of mice that vary in sensitivity to malaria infection^12^. We measured weight, temperature, red blood cell (RBC) concentration, parasitemia (proportion of infected RBC per total RBC), and parasite density (millions of parasites per microliter of blood). We examined mice on day 0, as an internal uninfected control for each mouse, and on days 5 through 15, as peak pathology occurs during this interval. To determine the range of phenotypes resulting from *P. chabaudi* infections, we tested the 8 founder strains of the collaborative cross to measure the limits and timing of disease phenotypes^10^.

As a first approach to measuring immune diversity, we plotted time series spaghetti plots for each phenotype and found a broad range of results (Figure 1A-D): Some mouse strains showed high pathology when infected under this regime while others suffered essentially no disease. For example, the WSB/EiJ mice did not lose weight, become hypothermic, or become severely anemic when infected with *P.chabaudi* AJ, and their infection peak occurred later than the other mice. C57BL/6 mice had mid-range phenotypes with about half of the other mouse strains doing better or worse. We found that the DO mouse disease outcomes largely recapitulate the spectrum of parental strain disease responses though some areas of disease space are occupied by the DOs and not the parents.

**Figure 1.**
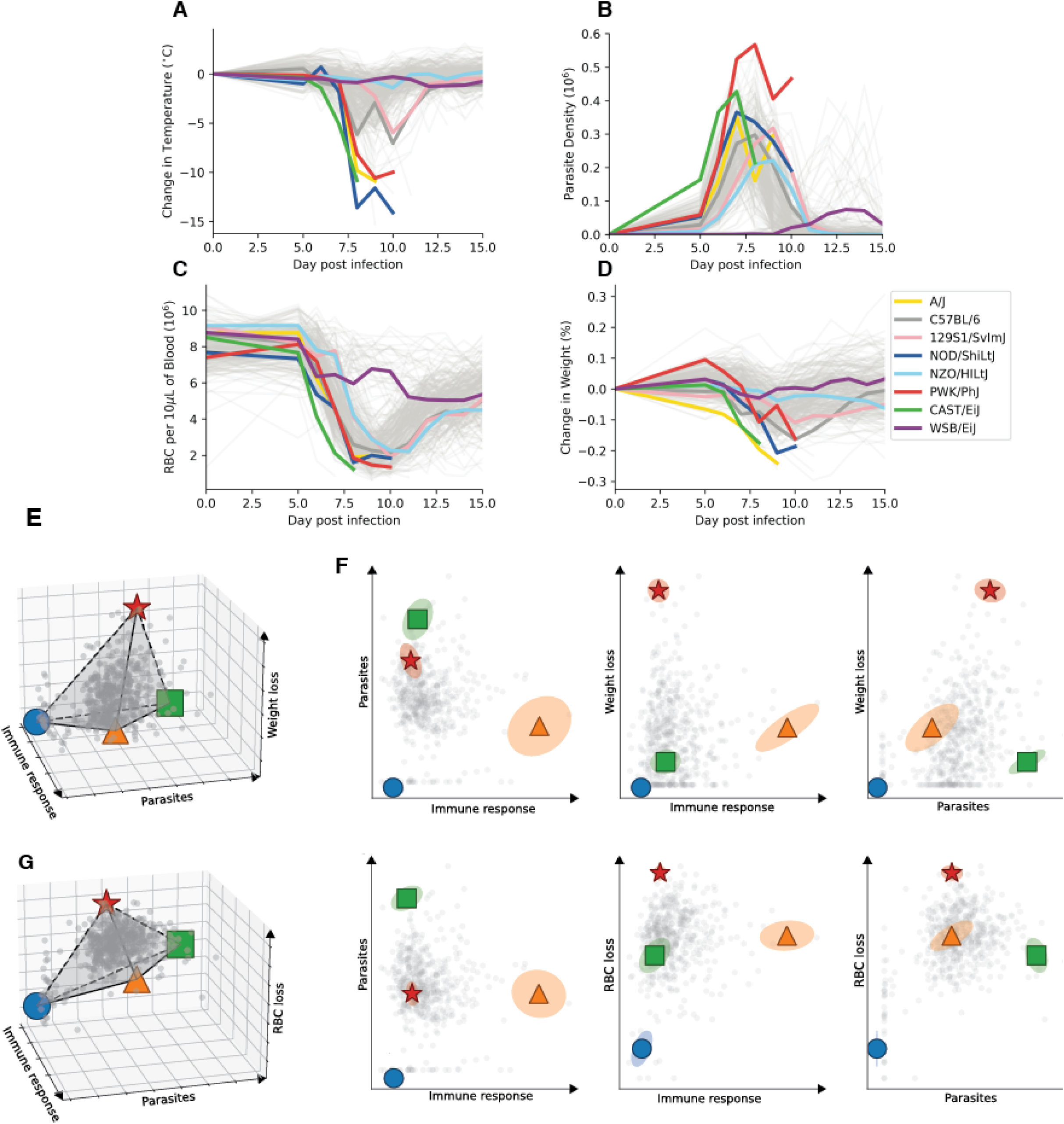
The infection archetypes resulting from malaria infection in diverse mice. Spaghetti plots showing the time series outcomes for P.chabaudi infected DO mice and the eight parents of the collaborative cross: A. Temperature loss (sick animals become cool). B. Parasite Density. C. Red blood cell loss. D. Weight loss. E. Three-dimensional view of the DO and parent mice in microbes by parasite clearance rate by weight loss. This is plotted using Z scored values for the parameters. F. Three two-dimensional views of microbes by clearance rate by weight loss showing the positions of the four archetypes and the uncertainty of those positions (see methods) Parameters are reported as Z scores. G. Three-dimensional view of the DO and parent mice in microbes by parasite clearance rate by anemia. H. Three two-dimensional views of microbes by clearance rate by anemia show the positions of the four archetypes and the uncertainty of those positions. Parameters are reported as Z scores.

To understand how the phenotypes relate to each other we summarized the course of infection as we might when plotting disease tolerance. We picked the largest decrease in health from the baseline for each mouse and plotted these summaries against each other. We plotted the summary of the disease course of each individual on a 3D space consisting of maximal parasitemia, maximal damage, and maximal realized immune response (calculated indirectly for each mouse using parasite burst size, as described in Metcalf et al. 2012^13^). The dataset was found to have roughly a tetrahedral shape, with 4 archetypes (Figure 1 E, F). There was one archetype (red star) that became ill and three other archetypes (blue circle, green square, orange triangle) that remained healthy.

The three parameters we measured for damage (weight loss, temperature drop, and anemia) yielded 2 types of distributions in phase space (Figure 1 E-G, Supplemental Figure 1). In the first pattern, which is seen with weight and temperature, phenotypes at a given parasite load varied from perfectly healthy to some level of pathology. The peak damage point (red star) was found on the high microbe side of the graph. Anemia phenotypes varied from a moderate level of pathology to severe. Here the peak damage point (red star) was shifted away with respect to those seen with weight and temperature.

To better understand the characteristics of each archetype, we developed a simple mathematical model to describe the dynamics of the microbe, the host immune response, and the host damage. The model is general and applies not only to malaria but to a wide variety of pathogens.

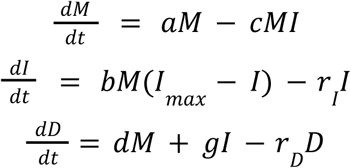

*M* stands for the microbe’s amount; *I* stands for the host’s immune response; and *D* stands for host damage. The variables a and c are the microbe growth and clearance rates respectively. The variable b is the rate of immune induction. The variable *b* is the *I*downregulation rate for the immune response. The variables d, g, and *r*_*D*_ are the damage rates driven by microbes and the immune response, and the repair rate. There are three general scenarios for damage (two are depicted schematically in Figure 2): In the first, only the microbes cause damage; in that case, *g* = 0, *d* ≠ 0. In the second case, only the immune response causes damage; in that case, *d* = 0, *g* ≠ 0 . The third case is when both immune response and microbe activity cause damage; in that case, *d* ≠ 0, *g* ≠ 0. To emulate the scenario in the experiment, we randomize parameters for 200 realizations of the model, and let it evolve according to the dynamical equations above for each of the first two scenarios. For each individual, we summarize the maximal microbe level, immune response, and damage it reached along the simulation, and plot it on a 3D space, much like the experimental data. The scatter plot created a cloud similar in shape to that of the experimental data; it had a roughly tetrahedral shape, and archetypal analysis using PCHA (Principal Convex Hull Analysis)^14^ found 4 archetypes in relative locations similar to those of the data.

**Figure 2.**
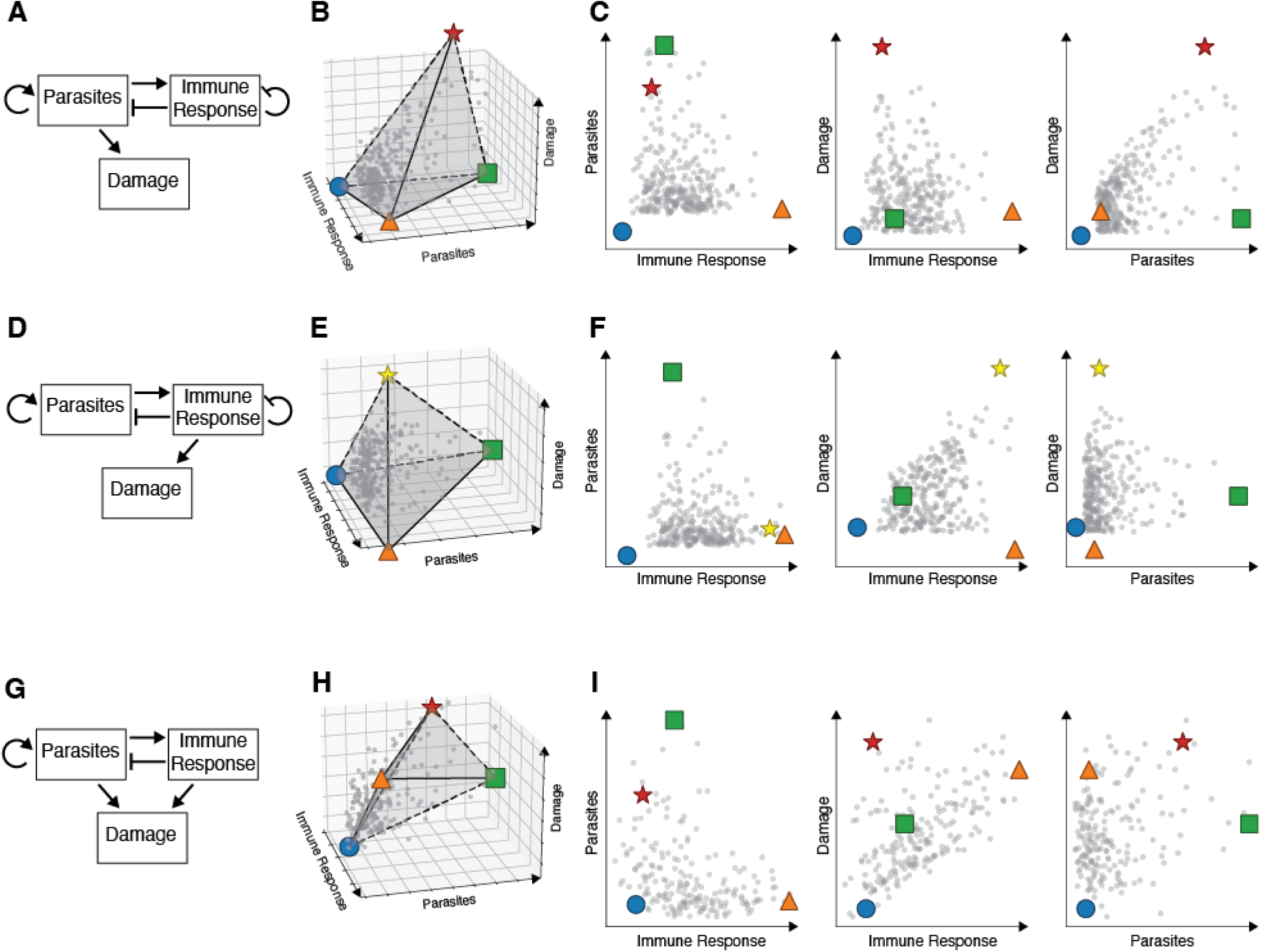
Mathematical model of disease space distribution. The box and arrow diagrams in A, D, and G show forms of the model where damage is either entirely microbial-mediated, entirely immune-mediated, or a combination. B, E, and H show the respective 3D spread of the sampled points and archetypes. C, F, and I show the respective projections on each of the 2D spaces: MxI, DxI, DxM. The four archetypes are marked with the same code as used in Figure 1. Parameters are reported as Z scores.

Another shape that can result from this model resembles the shape of the data cloud in the anemia case (Fig 1G, H). Such a shape is produced when there is both immune- and microbe-mediated damage (*d*≠0, *g* ≠ 0). In that case, non-zero levels of damage are observed in both low immunity (microbe-mediated damage) and low microbe levels (immune-mediated damage). Since the RBC level is very sensitive both to the parasite and to the immune response, sample ranges for the two parameters have a relatively high lower bound, resulting in the lifting of the M-I+D+ relative to the D axis.

In our model, the microbial damage and immune damage scenario share three archetypes that remain healthy and don’t vary with the source of the damage: (M↓, I↓, D↓) (blue circle),(M↑,I↓,D↓) (green square) and (M↓,I↑,D↓) (orange triangle in both fig1 and 2). These form a triangular base in MxI space and together these define a plane of minimum possible damage. The archetype with maximal microbial damage is found at (M↑,I↓,D↑) (red star in Figure 2B,C) which sits roughly above the M↑I↓D↓ archetype. The archetype with maximal immune damage is found at (M↓, I↑, D↑) (yellow star in Figure 2 E, F) and sits roughly above the M↓,I↑,D↓ archetype. The shape of the space we observe using temperature and weight as endpoints resembles the microbial damage tetrahedron while the anemia space appears more like the immune damage model.

To explain the behavior of each archetype, we used task inference^9^ to sort the model’s rates (microbe growth and clearance, immune induction and quenching, microbe or immune-induced damage and damage repair) to reveal the underlying tasks in which each archetype excels. (Supplemental figures 2,3). We distinguish these archetypes based on their resistance and tolerance to the infection. Here we use “resistance” to describe hosts that can readily clear microbes and “tolerance” to describe hosts that have a relatively low ratio of damage/microbe load. The five archetypes found in microbe and immune-mediated damage models provide examples of all four possible combinations of resistance and tolerance. The high damage archetypes (yellow and red stars) are notable for their high rates of damage and their low repair rates while microbe growth rates, killing rates, and immune sensitivity or magnitude are not enriched. The microbe-driven damage (red star) archetype lacks both resistance and tolerance. The immune-driven (yellow star) archetype has high resistance but low tolerance. One archetype in the triangular base is a tolerant host that has high microbe loads but little damage (green square). This archetype has many microbes because, like the red star archetype, its immune response isn’t particularly sensitive, but the host doesn’t suffer the microbial damage the red star suffers. The second healthy archetype is both a resistant and tolerant host that has the highest immune response, yet has a low immune damage rate (orange triangle). The last archetype is resistant and has low microbes, low immunity, and low damage (blue circle). This archetype is characterized by changes in the growth rate of the microbe and the efficiency of immune killing. It provides a different type of resistance as compared to the orange triangle, which has a large but only moderately effective immune response.

The rate constants of the model provide a sort of “health weathervane;” instead of showing the wind direction, these health vectors predict changes along three axes (Figure 3). Damage induction and repair are the easiest vectors to understand. They move a host up and down according to the damage scale. The microbe killing vector extends out the M-I-D- (blue circle) archetype and growth points in the opposite direction. The vector describing the sensitivity of the immune response to induction is directed out of the M-I+D- (orange triangle) archetype when damage is driven by microbes. When damage is driven by the immune response, this vector is rotated ninety degrees to exit through the M-I+D+ (yellow star) archetype.

**Figure 3.**
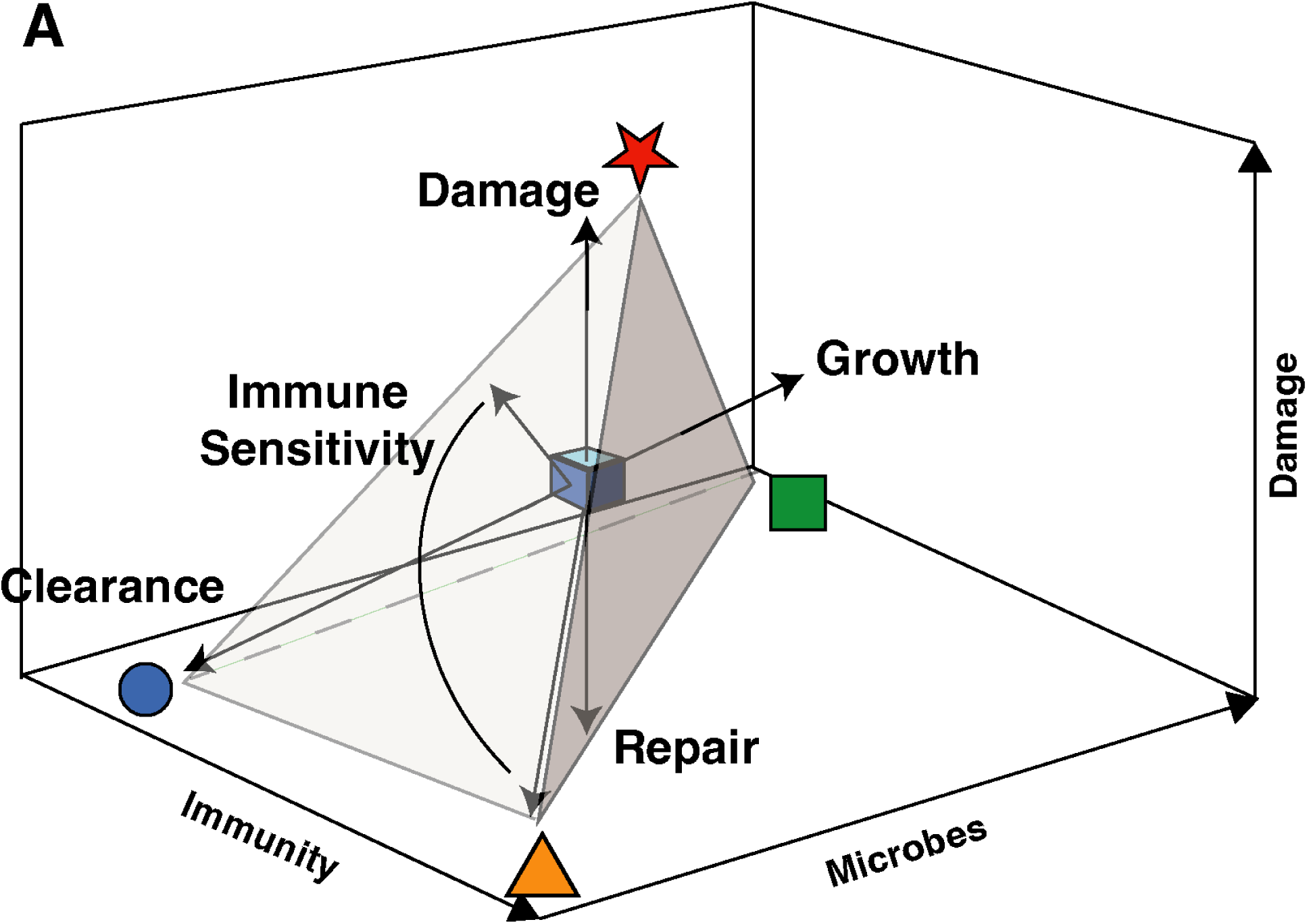
Vectors controlling the position of hosts in disease space. The tetrahedron shows the distribution of host phenotypes in MID space. The blue cube represents a host in the middle of this data cloud and the vectors show how this point will move when different rate constants are changed. Damage and repair move opposite each other along the same line. The same is true for growth and clearance. Immune sensitivity produces different vectors depending upon how much damage is caused by the immune response, as shown by the arc.

Our experimental data and mathematical model suggest the existence of multiple infectious disease archetypes. We reasoned the archetypes that do poorly with malaria might have been selected by host-microbe interactions where another archetype is favored. To test this idea, we examined the published outcomes of diverse mice infected with TB, IAV, and SARS-COV (Figure 4)^15–17^. We can’t compare the behavior of DO mice between these experiments as these mice are individuals and can’t be retested or regenerated; however, we can compare the results of the parents of the collaborative cross, which were also tested in these studies. We took a MID plot for a malaria infection and colored the parent mice with respect to their response to other infections (Figure 3). In the case of both TB and IAV, the mice that show high parasite loads in malaria infections are resistant to TB and IAV. The reverse is also true – WSB mice are resistant to Plasmodium but not resistant to TB and IAV. The SARS-COV-1 infected mice were healthiest in the middle of the malaria data, suggesting that there could be an archetype that would become visible if we found an appropriate third dimension. The WSB mice do not suppress the growth of TB, IAV, or SARS-COV-1, suggesting that if these mice raise immune response, it is inappropriate for those infections.

**Figure 4.**
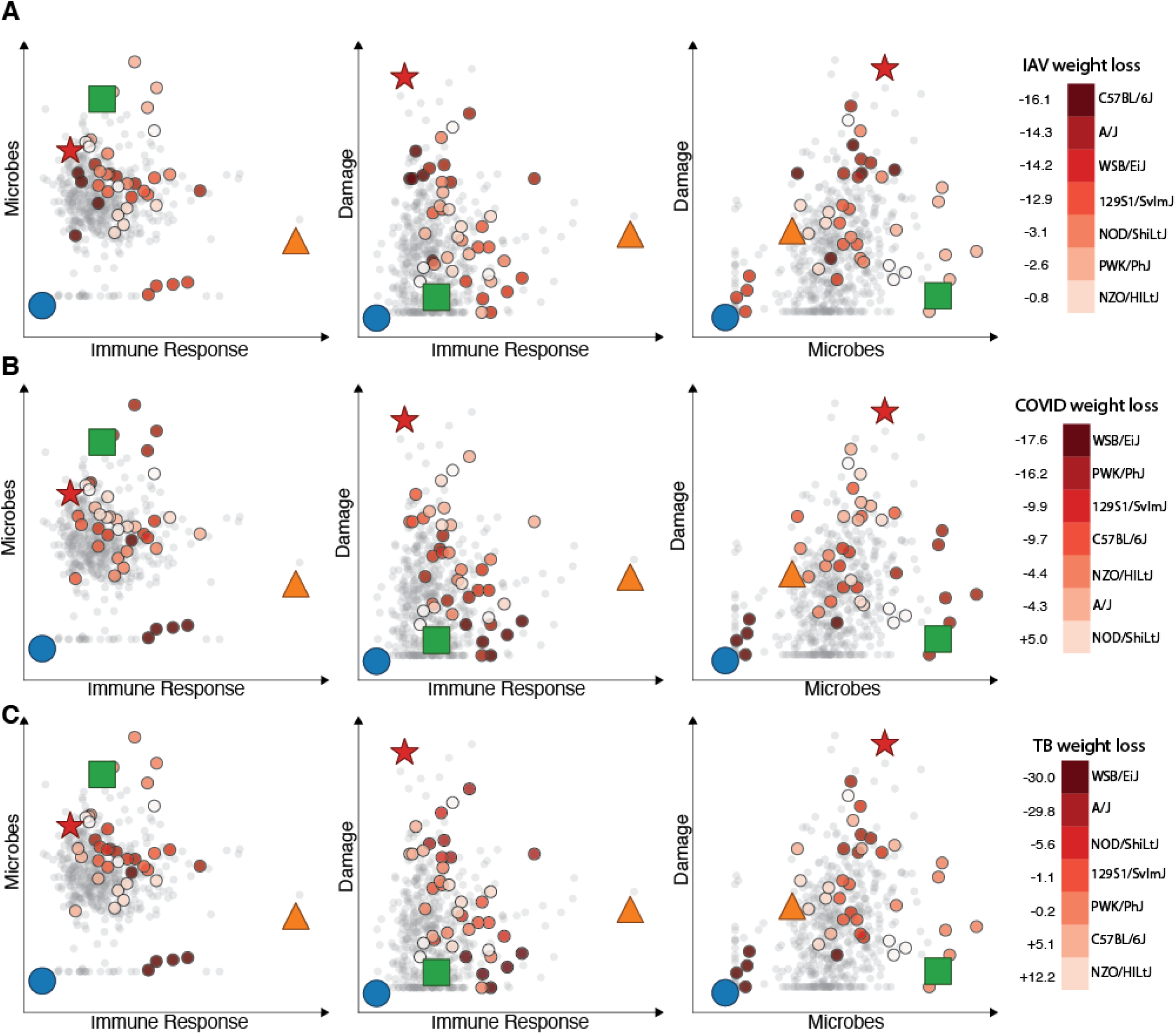
Distribution of other infectious disease outcomes in malaria disease space. Each of A, B, and C shows three 2D views of disease outcomes from the malaria infections. The colors represent weight loss results for the parent strains as reported for: A - IAV, B - SARS-COV1, and C – TB. Three-dimensional views are shown in Supplemental Figure 4. Parameters are reported as Z scores.

## Discussion

Organizing immunological space according to archetypes will help us find hidden archetypes, predict changes in responses to different pathogens and better compare our model systems to real patients. For example, the diversity of responses we see from mice as a species shows us that there isn’t one way that “The Mouse” deals with a given infection, just as there is no one way humans react^18^; rather there are five archetypical responses, everything in between, but nothing outside of this range. The shape of this space is something that should be universal and will hold across species that regulate microbe load with an immune response and suffer damage. We anticipate this architecture will exist at different scales, ranging from cells to species, so long as we are describing microbe load, the immune response and damage.

Our work links three schemas for discussing infections by showing they are three different views of the same system: inflammation regulation, the damage response framework, and disease tolerance. Understanding how microbial products can drive inflammation teaches us about mechanism, but doesn’t consider health, which drives the evolution of immunity. The damage response framework connects health to immunity but doesn’t address the immune response’s ability to control the microbes driving the infection^4,5^. Disease tolerance looks at the beginning and end of the infection process but leaves the middle as a black box^2,3^. If we consider that a host will occupy a pentahedral space, then each of these 2D views conflates two pairs of archetypes because of overlapping data points. These confused archetypes are resolved by adding back the missing dimension.

The idea behind Pareto optimization is that archetypes are ideal states for dealing with particular stresses. What are the stresses that match the 5 archetypes we observe in our model? The answers lie in supplementary Figures 2 and 3. The archetype at M↓I↓D↓ (blue circle) may be due to changes in the behavior of the microbes. Our model shows that the pathogens in these animals have increased sensitivity to antimicrobials and are slower growing. Either of these rates could be controlled by the host or pathogen, but the only source of variation in our experiments was the host. This suggests the hosts in this archetype are responsible (directly or indirectly) for the changes in microbe growth rates and antimicrobial sensitivity. For this reason, we call this archetype “Anansi” after the African trickster god^19^, as the microbes may have been misled into growing slowly and becoming more sensitive to the immune response. This is an effective way of clearing malaria parasites.

The M↑I↓D↑ (red star) is very sensitive to microbial damage, repairs poorly and it has a slow rate of immune induction. Again, these rates could be due to host or microbial factors, but we only varied the host in our experiments. With regards to the immune sensitivity, these hosts could be either blind to pathogens or raise the wrong immune response. We think that both of these things happen as this archetype contains different strains of mice, some of which are good at fighting other pathogens, and others that do not do well. We refer to this archetype as “Ate,” for the Greek goddess of mischief, folly and ruin; the host has made an error in selecting the immune response and must suffer the consequences. We presume that this archetype does poorly with malaria because of a trade off in its ability to fight other pathogens.

The M↑I↓D↓ (green square) archetype resembles the M↑I↓D↑ archetype with the exception that it is tolerant of microbes because it prevents damage from occurring. This could either be through manipulation of the microbe or by blocking the damage. This tolerant archetype is the position in disease space that could accommodate mutualism and commensalism. This archetype also provides a home for healthy super-shedders. We call this the “Honey Badger” archetype, after the internet meme describing the tolerance of honey badgers to distress. Such an archetype could be selected in an environment where the immune response cost outweighs the damage, is too damaging on its own or the relationship is beneficial.

The M↓I↑D↑ (yellow star) archetype suffers self-harm caused by a functional immune response. This might be a necessary tradeoff to survive extremely pathogenic organisms, but appears as an over-reaction when facing more mild pathogens. We refer to this as the “Wolverine” archetype, named after the Marvel superhero who wounds himself every time he prepares for battle^21^.

The M↓I↑D↓ (orange triangle) archetype has a rapidly induced immune response and is also good at preventing and repairing damage. We call this the archetype “Paladin” after the Dungeon and Dragons character that can both fight and heal^22^. This along with the Anansi archetypes seem to be the best solution for clearing malaria parasites.

Three of the five archetypes are relatively successful at clearing pathogens and use two different approaches to do this, according to our model. The Anansi hosts appear successful at limiting pathogens by decreasing the microbial growth rate and increasing microbial sensitivity to antimicrobials. In contrast, the Paladin and Wolverine hosts clear parasites by having sensitive immune responses that turn on quickly.

Three of the five archetypes are relatively healthy infection outcomes and highlight that microbial clearance isn’t the only way to decrease the impact of infections. By understanding the starting position of an infection in disease space and the vectors we can apply to the host, we could become better at choosing treatments. Treatments that lower immune sensitivity, like cortisol or anti-inflammatories can reduce self-harm and would be useful for helping both Wolverine and Ate type infections. The treatments are riskier when hosts face damage from increased pathogen loads in Wolverines.

Antimicrobials will push hosts towards the Anansi archetype and will be effective in Ate type infections which do not produce a strong immune response. Antimicrobials will be less useful in Wolverines, who produce an effective microbe killing response. Damage reduction drugs like analgesics for pain or anti-oxidants for reactive oxygen damage will push hosts down towards the MxI plane of the graph. The same is true for damage repair treatments like oxygen for respiratory distress or rehydration therapy for cholera.

In the same way that a species of host can vary through a pentahedral space, we expect that different species of hosts will be distributed in a similar manner. Zoonoses can have a carrier host that is the Honey Badger archetype and we notice the microbe because it causes Wolverine or Ate disease in humans. Examples of reservoir hosts that suffer little damage from their microbial passengers occur across the animal kingdom and include deer mouse (sin nombre virus)^23^, Fruit bats (nipah virus)^24^, Egyptian rousette bat (marburg virus)^25^. This leaves us wondering about the nature of the difference between humans and these carriers, but in doing so we risk falling into a survivorship bias trap. Animals that have the Anansi and Paladin archetypes would also be interesting to study as they must have mechanisms for limiting disease, but we miss them in nature precisely because they do not suffer from disease. These healthy animals in addition to the Honey Badger archetype could help teach us how to avoid pathology.

We anticipate that the pentahedron revealed in MID space will be found not only in animals of one species but at different scales. We expect to see shapes like this ranging from different animal species down to the behavior of single cells. Accordingly, it is difficult to imagine one infectious disease archetype providing a good solution to all organs in the body and we might expect different organs to be distributed in a pentahedral manner in MID space. This is an extension of the idea that disease tolerance will vary between tissues^26^. For example, intense fighting archetypes could work well in tissues that can tolerate damage; however, tissues that don’t regenerate will require less damaging responses or an insensitivity to damage. We anticipate that different parts of the body will be tuned to different archetypes and this could create tradeoffs between organ systems in terms of their sensitivity to various pathogens. Our studies already provide some evidence of intra-host outcome variation. We followed three different health outputs and each of these systems has its own tuning with respect to parasite load. Human malaria provides additional evidence of intra-host variation of organs ^27^. Malaria causes different syndromes in humans, differentially harming the brain, lungs, liver, kidneys, gut, placenta, and hematopoiesis. We anticipate that there will be an overall immune tone set for each individual and that their organs will vary within limits with respect to this set point.

In our experiments, we used disease outcomes that summarized the infection, which is simple to do in the lab, but harder to accomplish in nature or the clinic. It would be useful to develop a series of stress tests that measured key immune, damage production, and repair rates. Medicine already has cardiac, pulmonary, glucose, and kidney stress tests. Inflammatory stress and damage recovery tests would be a useful addition to our assays of health. This would help us measure the rate constants that lead to different archetypes. It may be possible to predict how a patient will behave by taking a personal natural history approach. If we recorded severe reactions to identified childhood infections to place a patient on the map of disease space, this would help predict future responses.

## Methods

### Mice

Female mice were purchased from The Jackson Laboratory (JAX) (Bar Harbor, Maine, USA) and maintained in the Stanford University animal facility. Mouse strains used in this study are listed in Table S1. All experiments were performed in accordance with institutional guidelines and approved protocol (APLAC-30923). The first batch of DO mice was from Generation 23, Litter 2. The last batch of DO mice was from Generation 30, Litter 1. Mice of each parental strain were included in experiments for genotyping controls and infection phenotype analysis (N=4-7 for each parental strain).

All mice were acclimated in cohoused cages of 5 animals in the Stanford University animal facility for 7–10 days prior to being used in experiments. Female mice of a similar age (8-15 weeks old) were infected with *Plasmodium chabaudi chabaudi* AJ (Malaria Research and Reference Reagent Resource Center [MR4]) to limit variance in the murine response to this parasite caused by sexual dimorphism and age. The DO mice were not shuffled to randomize microbiota on arrival to the facility due to concerns of aggression towards unfamiliar cagemates.

### Temperature Probes

Mice were anesthetized locally with 2% lidocaine solution (100 μg delivered per dose) and implanted subcutaneously with electronic temperature and ID transponders (IPTT-300 transponders, Bio Medic Data System, Inc) one week prior to *P. chabaudi* infection. Temperatures were recorded using a DAS-7006/7s reader (Bio Medic Data System, Inc).

### Infection

C57BL/6 female mice were given intraperitoneal injections of 100 μL of thawed *P. chabaudi chabaudi* AJ-infected RBCs (iRBCs) from the parasite stock described previously. In order to monitor parasitemia, thin blood smears were prepared from tail blood, methanol fixed, and Giemsa stained (Gibco-KaryoMAX). Parasitemia was quantified under light microscopy at 100x magnification daily until parasitemia reached 10-20% (9 days post-infection). Then experimental mice were injected intraperitoneally with 100 μL containing 10^5 freshly obtained iRBCs diluted in Krebs saline with glucose.

### Longitudinal infection monitoring

Weight and temperature were recorded daily between 9 AM and 12 PM during the experiment. In situations where mice were not implanted with temperature probes, the temperature was determined using a rectal thermometer (BAT-12, Physitemp Instruments Inc.). Approximately 19 μL of blood was collected daily via tail nicking of restrained mice using sterilized surgical scissors. Sodium acetate warmers were used on occasion to increase tail blood flow. Blood was collected into EDTA-coated 50 μL capillary tubes to inhibit clotting. Beginning on day five, roughly 2 μL of blood per animal was used to generate thin blood smears, which were subsequently prepared and counted as detailed above to measure parasitemia. Each day, 2 μL of blood was diluted in one mL of Hanks’ Balanced Salt Solution (HBSS) to count the number of RBCs.

Absolute RBC counts were obtained on an Accuri C6 Flow Cytometer (BD Biosciences) using forward and side scatter.

Parasite density (parasitized RBC per volume of blood) = infected RBC total RBCAbsolute RBC count

#### 2. Estimating Immune Response

The immune response was quantified using the method developed by Metcalf. et al.^13^, which integrates RBC count and parasitemia to compute an immune activity score.

Specifically, the immune response was approximated by the *bystander killing* each day, calculated by the difference between the susceptible (uninfected) RBCs of each day and the parasitemia and uninfected cells from the previous day. The calculated immune response values were utilized for further analysis.

#### 3. Extraction of Maximal Values

For each mouse, the maximal values of three key variables were extracted from the time series data:

● **Maximal parasitemia**: Defined as the highest recorded parasitemia value during the experiment.
● **Maximal damage**: Estimated using the maximum absolute value of RBC count difference from day 0, the maximum absolute value of weight change from day 0, and the maximum absolute value of temperature from day 0.
● **Maximal immune response**: Determined using the highest computed immune response score from the method described above.

#### 4. Normalization and Comparison Across Mice

To compare inter-mouse variation, all maximal values were standardized using z-scores, calculated as:

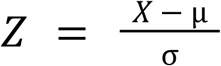

where *X* is the individual maximum value, µ is the mean across all mice, and σ is the standard deviation.

#### 5. Model Description

A three-dimensional system of ordinary differential equations (ODEs) was used to describe the interactions between pathogen load, immune response, and damage. The system, with its variables and parameters, was discussed in the results. The ODEs were solved numerically using the Euler integration method.

#### 6. Parameter Sampling and Simulations

Many model parameters could not be directly estimated due to their abstract nature or the difficulty of obtaining precise measurements. Instead, parameters were selected to ensure that the simulated variables remained within biologically plausible ranges.

Parameters were drawn from the following distributions:

● **Microbe growth rate (a):** Daily differences in parasitemia levels were calculated for each mouse. The average positive (growth) values were used to determine the distribution parameters. The model parameter values for microbe growth were then sampled from a Gaussian distribution with the same mean and standard deviation as the empirical growth rate.
● **Maximal immune response (I_max):** The maximal level of the logarithm of the immune response was plotted, and the model parameter for I_max was sampled from a Gaussian distribution with the same mean and standard deviation as the observed data.
● **Damage repair rate (rD):** Daily differences in damage indicators (weight and temperature) were computed. Similar to the microbe growth rate, the average of the positive differences was used to define the distribution, and values for rD were sampled from a Gaussian distribution with the same mean and standard deviation.

The parameter ranges and distributions can be found in table 1. Adjustment for RBC variation is found in table 2 (all parameters that are not mentioned in table 2 were set to be drawn from the same distributions as in table 1)..

**Table 1:** Parameter values.

| Parameter | Symbol | Range |
| --- | --- | --- |
| Microbe growth rate | a | $a \in N(0.9, 0.3)$ |
| Immune stimulation | b | $b \in U(1, 7)$ |
| Microbe killing | $c$ | $c \in U(2, 5)$ |
| Microbe-mediated damage | $d$ | $d \in U(0, 5)$ |
| Immune-mediated damage | $g$ | $g \in U(0, 5)$ |
| Damage repair rate | $rD$ | $rD \in N(1, 0.3)$ |
| Immune shutdown rate | $rl$ | 0.1 |

**Table 2:** Parameter values in RBC variation.

| Parameter | Symbol | Range |
| --- | --- | --- |
| Microbe-mediated damage | $d$ | $d \in U(2, 5)$ |
| Immune-mediated damage | $g$ | 0.1 |

To validate this approach, we compared simulation outputs to experimentally observed values, adjusting parameter distributions as necessary to maintain consistency with known biological constraints.

Each simulation was run for a total time of 5 days, with initial conditions set to I=0.001, M=0.1, D=0. The initial parasitemia level was so chosen because lower levels of parasitemia were not detected in the experiment. A very low, though nonzero, initial value of immunity level was chosen to prevent numerical artifacts. A total of 300 simulations were conducted. Simulations that reached damage levels higher than 3 standard deviations from the cohort average were disregarded. The total numbers of simulations used for the next steps were 217-296.

#### 7. Archetype Analysis Using PCHA

To characterize the behavior archetypes in the dataset, we applied the Principal Convex Hull Approximation (PCHA) algorithm^14^. The number of archetypes was set to 4 determined a priori, using δ=0.1. The PCHA algorithm was implemented using the Python package PyPCHA.

To estimate the error in the archetype location, we conducted 50 bootstrapping iterations, each sampling 200 points of the data. For each bootstrap iteration, we used the PCHA algorithm to find four archetypes. Then, the archetype locations were clustered into four, and the center of the cluster was interpreted as the final archetype location. The uncertainties in the archetype locations are represented in Figure 1 as ellipses, whose principle axes are determined by the covariance matrix of each cluster.

## Acknowledgements

We thank Rebecca Gellman, Elysse Grossi-Soyster, Carolina Tropini, Avi Mayo and Elizabeth Vaisbourd for reading the manuscript.

This work was supported by the following funding sources: Defense Advanced Research Projects Agency W911NF-16-0052 (D.S.S.) and a Stanford Discovery Grant (D.S.S.). Y.L. is supported by the Azrieli Foundation Graduate Studies Program.

Funders were not involved in study design, data collection, analysis, or interpretation, writing of the manuscript or decision on where to submit for publication.

---

**Supplemental Figure 1:**
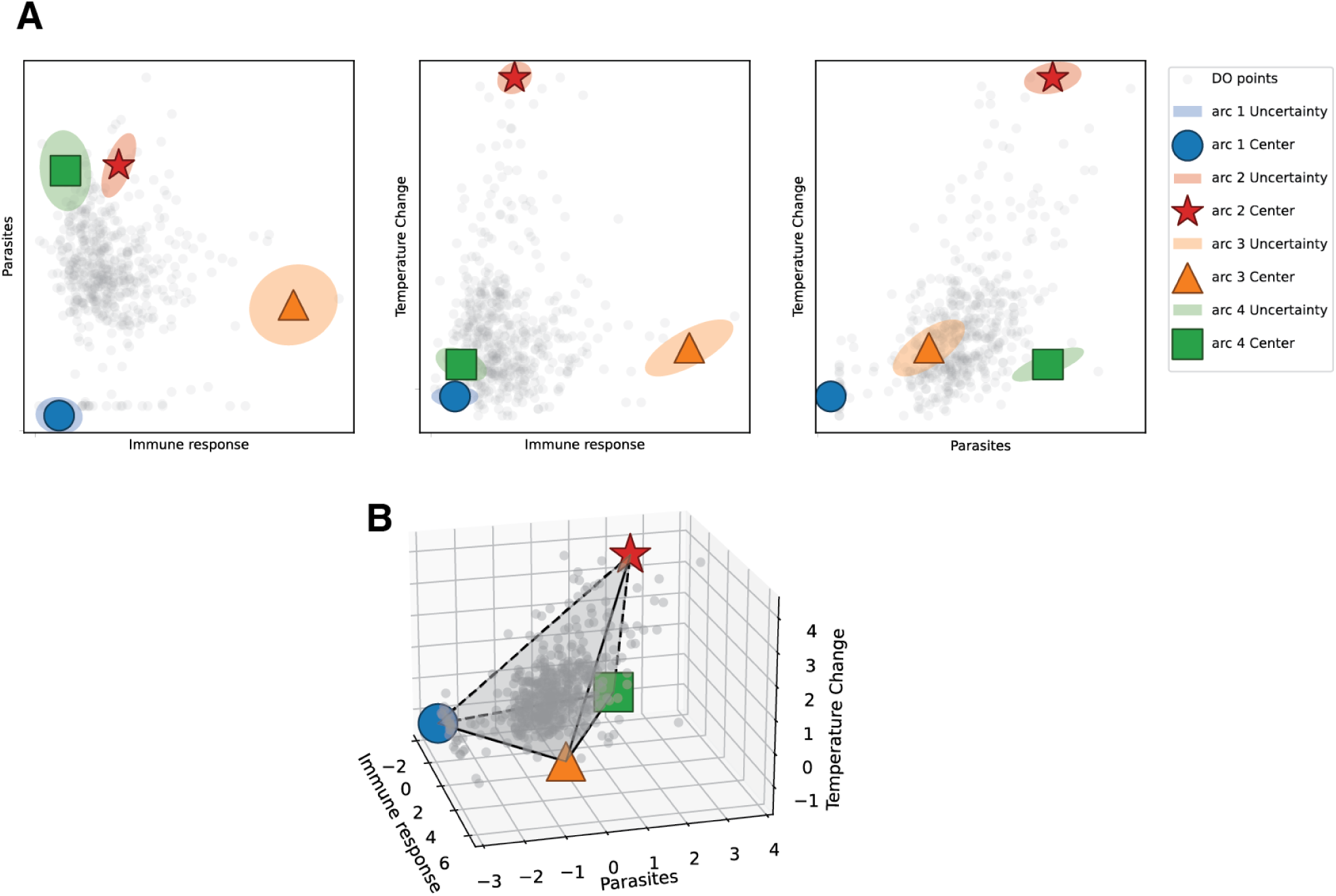
Archetypes for temperature change

**Supplemental Figure 2:**
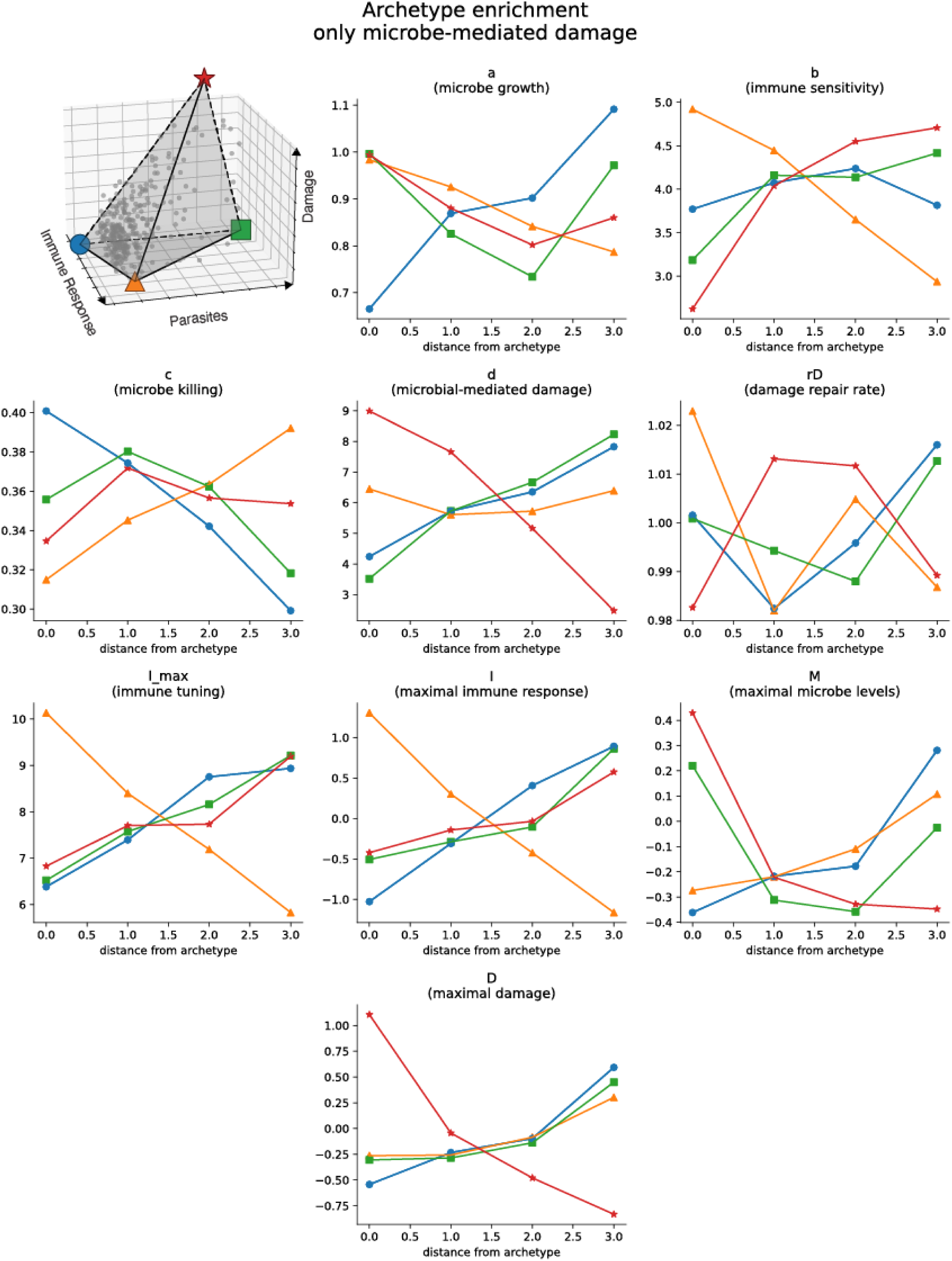
Archetype enrichment results for the microbe damage model.

**Supplemental Figure 3.**
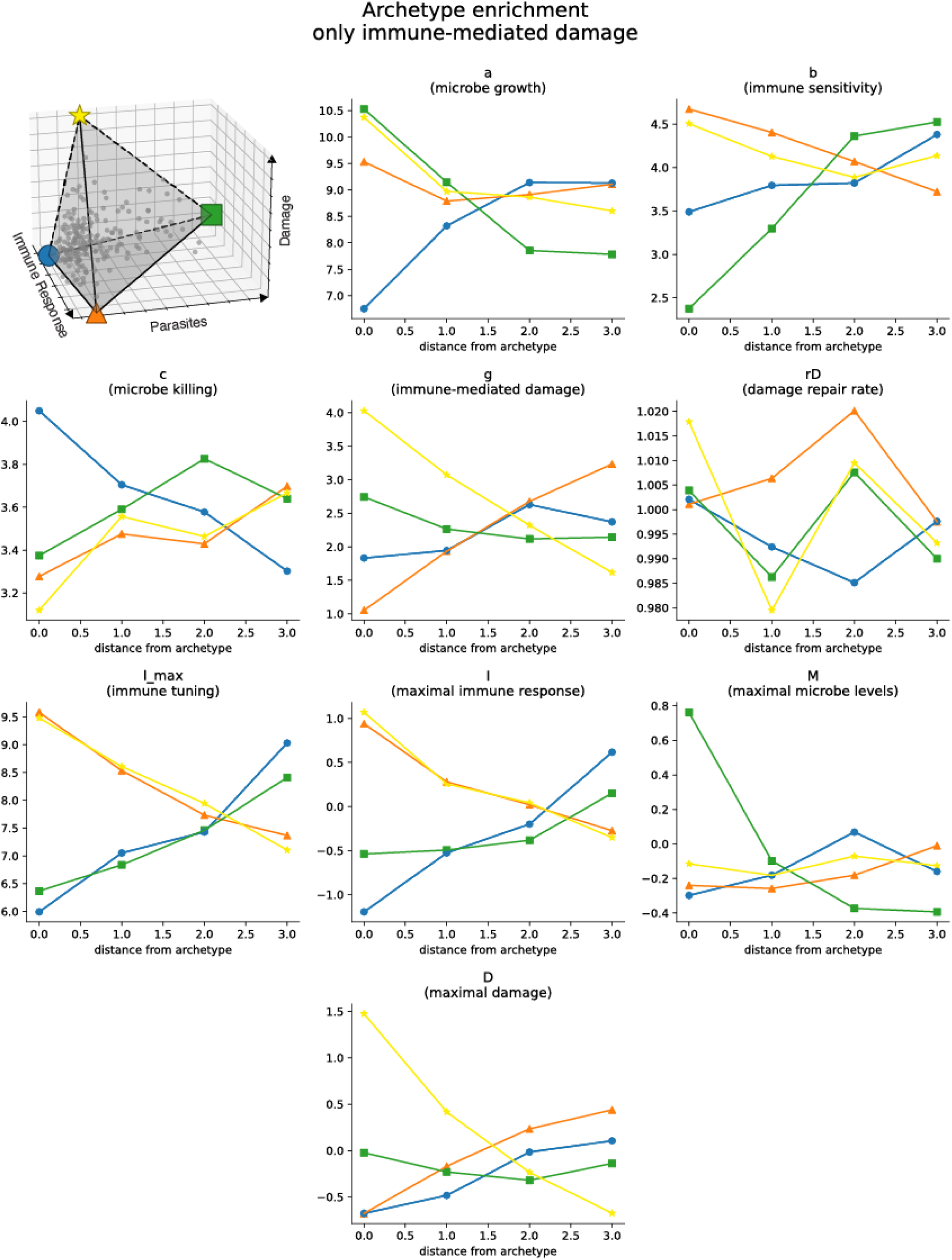
Archetype enrichment results for the immune damage model.

**Supplemental Figure 4:**
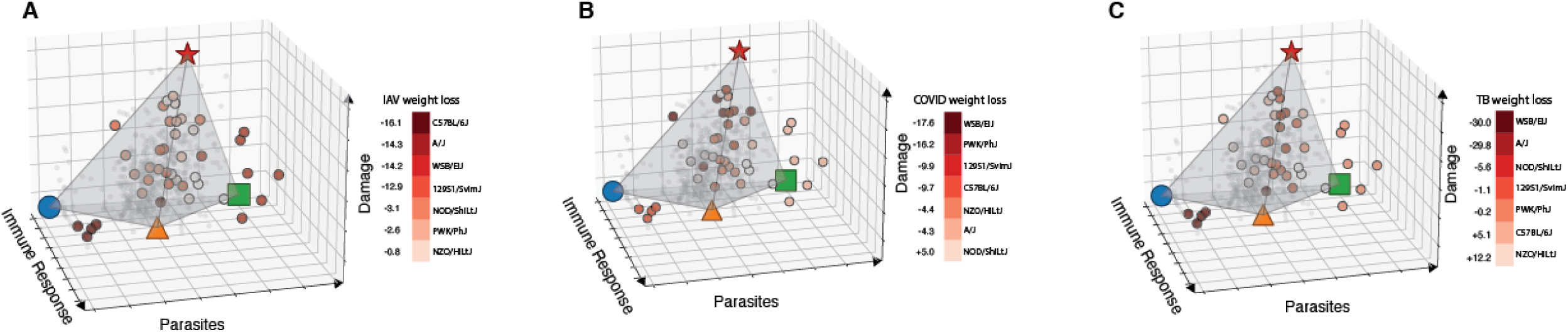
3D visualizations of the distribution of other infectious disease outcomes in malaria disease space.

**Supplemental Figure 5:**
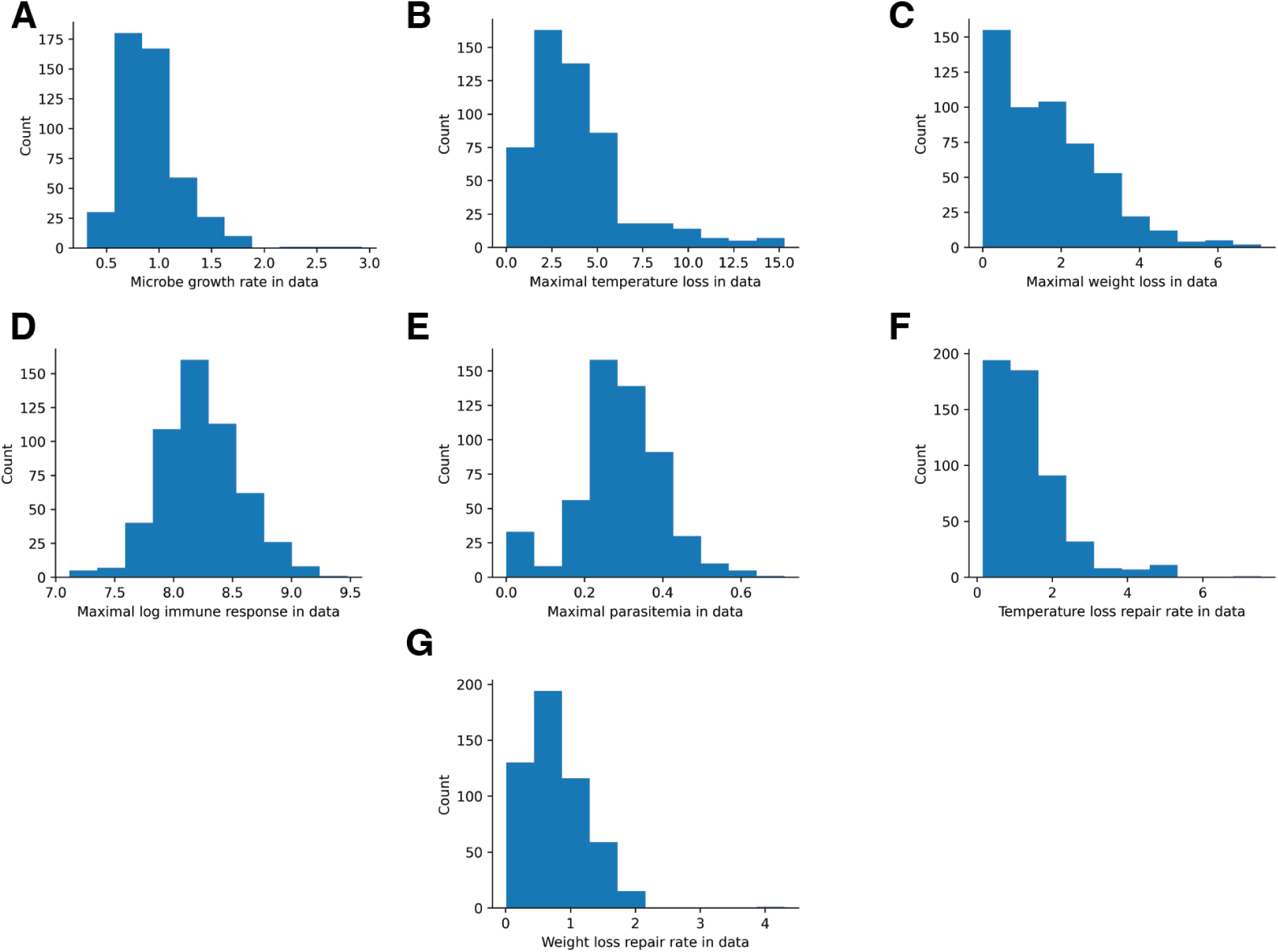
Distribution of calculated parameters in sampled mice.

